# Japanese macaques (*Macaca fuscata*) adjust inter-individual distance based on aggressiveness rather than rank differences

**DOI:** 10.1101/2025.07.21.664030

**Authors:** Kosho Katayama, Kazunori Yamada, Noriko Katsu

## Abstract

Group-living animals form societies structured around linear dominance hierarchies. However, social relationships within groups range from relaxed to tense, with some subordinates staying proximity of dominants, while others fleeing at their mere sight. Despite this variation, the effects of personality in shaping these relationships remains unclear. In this study, we investigated whether subordinates adjust their flight initiation distance (FID) according to the dominant’s aggressiveness. We collected video recordings of supplanting behaviours between non-kin dyads in Japanese macaques (*Macaca fuscata*). For each dyad, we evaluated these supplantings whether the subordinate’s FID was greater or less than two meters. To assess aggressiveness, we conducted 20-min focal observations of 18 females, recording of threats and aggressions. We calculated repeatability of aggressiveness as a measure of dominant’s personality. Rank differences were determined based on supplantings and aggressive interactions. As a result, aggressiveness showed high repeatability, independent of rank and sex. Subordinates were more likely to flee at greater distances (FID > 60 cm) when paired with more aggressive dominants. Rank difference did not significantly influence FID. Whether the dyads exhibiting FIDs were regular grooming partners did not significantly predict their FID. Our findings suggest that Japanese macaques adjust inter-individual distance not based on rank differences but in response to aggressiveness. These results highlight the importance of incorporating personalities into studies of primate social relationships.

## Introduction

A variety of group-living animals form dominance hierarchies. Individuals within these groups are capable of recognizing conspecifics and understanding their relative rank positions (Massen et al., 2014). Structuring social interactions based on these dominance relationships helps to reduce the risk of injury (Tibbetts et al., 2022). By understanding the rank of other individuals and adjusting their social behaviour accordingly, individuals can behave adaptively within their group.

Within primates, and specifically among macaques, there are variations in the steepness of the dominance hierarchies (Thierry et al., 2000). However, even in macaques with strict dominance hierarchies, there is a gradation in the degree of intimacy between individuals. These diverse relationships within groups have been actively explained by factors such as kinship and close relationships between non-kin individuals (Dunbar & Shultz, 2010; Silk et al., 2013). For example, subordinates frequently groom dominants (Nakamichi & Shizawa, 2003; Wu et al., 2018), thereby gaining tolerance, such as being permitted to forage (Oki & Maeda, 1973; Schino et al., 2007; Seyfarth, 1977). Although greater rank differences are associated with more intense aggression (Hemelrijk, 1999), grooming reduces the likelihood of being attacked by the recipient (Gumert & Ho, 2008). Particularly in groups with intense feeding competition, the greater the rank difference, the easier it is to obtain tolerance mediated by grooming (Barrett et al., 1999; Fruteau et al., 2009). This suggests that monkeys are aware of the rank differences among themselves and adjust their social relationships accordingly.

Furthermore, the variety of social relationships have been explained through the lens of affiliative relationships (Adiseshan et al., 2011; Schino & Aureli, 2009; Silk et al., 1999). Within groups, monkeys groom their kin more frequently than non-kin individuals (Sade, 1972; Silk et al., 2010). Although opportunities for aggression increase when individuals are in close proximity (Roubová et al., 2015), aggression remains low among kin despite frequent social interactions (Barrett et al., 2002; Silk et al., 2010). A positive correlation exists between the time or frequency of grooming given to a partner and the frequency of aggressive behaviour received from that partner (Carne et al., 2011; Schino et al., 2005; Silk, 1982). However, concerning the dominance relationships of macaque species, there is little research focusing on personality as a factor influencing these inter-individual relationships.

Consistent inter-individual differences among individuals within a conspecific group are called personality (Gosling, 2001; Réale et al., 2007; Sih et al., 2004a), and this has been shown to influence group relationships (McCowan et al., 2011). Individuals within a group might adjust their inter-individual relationships by perceiving each other’s personality, perhaps more so than by relying solely on their dominance relationships. For example, individuals within a group perceive not so much the ordinal rank of each other (e.g., 1st, 2nd, which reflects the hierarchical order of others), but rather a cardinal rank (the degree of an individual’s dominance expressed numerically by the ‘frequency’ of aggressive interactions). In a study on mandrills (*Mandrillus sphinx*), cardinal rank predicted grooming and avoidance behaviour better than ordinal rank (Schino & Lasio, 2019). Although there is little research verifying the perception of personality among animals within a group, if individuals perceive the cardinal rank of others, monkeys may understand each other’s personality of aggressiveness and adjust their social relationships accordingly. Empirical evidence on how monkeys recognize socially relevant personality traits in others may clarify why primates have personality. That is, personality may serve an adaptive function if its perception facilitates social adjustment and decision-making.

This study aimed to clarify whether Japanese macaques adjust their social relationships based on each other’s personalities. Specifically, the analysis focused on whether subordinates regulate the distance at which they initiate escape according to the aggressiveness of dominant individuals. Using observations of supplanting events, three hypotheses were tested within dominant–subordinate dyads. Hypothesis 1: FID increases as the rank difference between the individuals becomes larger. In other words, subordinates are expected to flee earlier from dominants who hold much higher positions in the ordinal hierarchy. Hypothesis 2: Dyads that share affiliative relationships, as indicated by grooming, tend to show shorter FIDs, even when their rank differences are large. Hypothesis 3: Independently of the effects of rank difference or grooming, the FID increases with the dominant individual’s baseline frequency of aggressive behavior. Subordinates likely escape earlier from dominants who consistently display higher levels of aggression. This pattern should hold even after controlling for rank differences and grooming interactions.

## Materials and Methods

### Ethics statement

All observations were conducted in accordance with the Regulations on Animal Experimentation at Osaka University, Japan, and were approved by the Animal Research Committee of the Graduate School of Human Sciences, Osaka University (No. R4-1-0). As the study site, Arashiyama Monkey Park, Kyoto, Japan, is privately owned, observations were made with the permission of the management.

### Study site and subjects

The study subjects were Japanese macaques from a free-ranging, provisioned group (Arashiyama group) living in Arashiyama Monkey Park (35°00′N, 135°67′E), Arashiyama, Kyoto city, Japan. Maternal kin-relations and individual ages in the group were recorded since the introduction of provisioning in 1950s (Fedigan & Asquith, 1991). The study was conducted from August 2023 to March 2025. In August 2023, the group comprised 126 individuals, including 6 adult (≥ 5 years) males, 59 adult females, 49 immatures (1–4 years), and 12 infants (<1 year) at the beginning of the study. In the Arashiyama group, breeding season was from September to March and the birth season was from April to August (Inoue & Takenaka, 2008). The subjects were 18 adult females (mean age ± *SD* = 19.06 ± 9.58 years; range: 5–31). The subjects were selected from adult females in the group, aiming for the widest possible age distribution and including only individuals with social ranks.

### Data collection

Behavioural observations were conducted between 09:00 and 16:30 h, when the group remained near the provisioning area. To record dyadic FIDs, video recordings (GoPro HERO8 Black, GoPro, Inc., California, United States and iPad 10th generation, California, United States) were made using ad libitum sampling method (Martin et al., 1993) in the vicinity of feeding areas. Subjects were also video-recorded during focal animal observations. Focal animal observation methods (Martin et al., 1993) were carried out in 20-min sessions to assess aggressiveness in dominant individuals within dyads. During each session, subjects were followed in a predetermined random order, and threat and aggression were recorded using all-occurrence sampling method (Table 1). The presence or absence of grooming interactions (1 = present, 0 = absent) between dyads in FIDs was determined from the focal observation data. The total observation time amounted to 173.03 hours for 18 subjects (mean ± *SD*: 9.61 ± 0.59 hours per individual). In addition to the video and focal sampling data, supplantings and aggressive interactions were recorded by the first author (K.K.) and the third author (N.K.) using the ad libitum sampling method to assess dominance ranks. These dominance rank interactions were recorded only when no other individuals were nearby, ensuring that the interaction involved only the two individuals directly concerned.

**Table 1.** Ethograms in FIDs from video recording, focal animal sampling, and ad libitum sampling method.

|  |  |
| --- | --- |
| *FIDs from video recordings* |  |
| long FID | In response to a dominant individual's approach, a subordinate runs or walks away before the inter-individual distance becomes less than or equal to arm's reach (60 cm). If a subordinate does not move away before the distance reaches 60 cm or less, they flinch or recoil. If a subordinate is engaged in feeding or grooming, the activity ceases. |
| short FID | When the inter-individual distance becomes less than or equal to arm's reach (60 cm), a subordinate runs or walks away to maintain a distance greater than 60 cm. In response to a dominant's approach, if the distance becomes 60 cm or less, a subordinate does not move away but flinch or recoil. If a subordinate is engaged in feeding or grooming, the activity ceases. |
| no response | A subordinate show no reaction and remain in place, when the distance is arm's reach (60 cm) or less. |
| *Focal animal sampling method* |  |
| rate threat | Hourly rate of threats directed at other individuals. Threats directed at person such as tourists, staffs, and an observer were also recorded. |
| *Focal animal sampling, and ad libitum sampling method* |  |
| rate aggression | Hourly rate of aggression such as biting, jumping on, and chasing other individuals. Approaching with threats directed at people such as tourists, staffs, and an observer were also recorded. |
| rate supplanting behavior/ receiving supplanting behavior | Hourly rate of moving another individual away from its original location when the subject approaches another individual without apparent agonistic behavior between two individuals. Or moving another individual away to avoid contact with the other individuals./ receiving supplanting behavior. |

### Data analyses

#### FIDs, rank differences, and aggressiveness of dominant individuals in dyads

From video recordings, we extracted a total of 96 dyadic supplantings using the video editing software Microsoft Clipchamp (version 4.2.10020.0). All 96 dyads were non-kin female pairs, included individuals over four years old, and allowed us to observe the dominant’s frequency of aggressive behaviors. For each dyad, we classified the FID as ‘long FID’ when the subordinate fled at a distance greater than arm’s reach (60 cm), ‘short FID’ when the subordinate fled at a distance less than or equal to 60 cm, and ‘no response’ when the subordinate did not flee, showed no reaction, and remained in place even when the approached by the other individual within 60 cm (Table 1). Two researchers with experience observing Japanese macaques coded the behavior from the video recordings of supplanting events. From the 96 dyads, we selected 47 in which their coding results matched for subsequent analysis.

Rank difference referred to the absolute difference in dominance rank between dyad members. Dominance ranks were assigned to 72 females aged 4 years or older by applying the Elo-rating method to 1,851 supplanting and aggressive interactions recorded through ad libitum and focal animal sampling. We selected this analysis rather than traditional matrix-based approaches, which reorder matrices by placing winners in rows and losers in columns to align interactions in the upper right. The Elo-rating method is superior to these approaches because it can account for situations in which dominant individuals are infrequently involved in supplanting events. All calculations used the ‘aniDom’ package in R (Sánchez-Tójar et al., 2018).

The analysis calculated the frequency of threat and aggressive behaviors by dominant individuals in FID dyads by dividing the number of such behaviors by the total number of focal observation hours for each individual. For females, the analysis excluded observation sessions during periods of estrus. Signs of estrus included the presence of a copulatory plug, facial reddening, or the emission of estrous calls. We calculated the repeatability of the aggression rate using the ‘rptR’ package (Stoffel et al., 2017) to examine whether aggression reflects personality.

#### Statistics

Before conducting the regression analysis, the analysis calculated the correlation coefficient between rank and aggression rate in 18 subjects to assess whether more dominant individuals engaged in more aggressive behavior. The analysis treated dyadic FID as a dependent variable for ‘long FID,’ for ‘short FID,’ and ‘no response.’ It applied a Generalized Linear Model (GLM) to examine the effects of the following explanatory variables: (1) the rank difference between dyads, (2) the frequency of aggressive behaviours exhibited by the dominants of dyads during focal observation, and (3) the presence or absence (1 = present, 0 = absent) of grooming interactions between dyads during focal observation. All analyses were conducted using R version 4.4.2.

## Results

### Number of FID dyads and rank of dominant subjects

Of the 47 dyads analysed, 94% (44 of 47) were recorded only once, and only three dyads were recorded twice (Table 2). The mean dominance rank of the subjects was 27.72 (*SD* = 19.84; range: 1–68). In the 47 dyads, each subject served as the dominant an average of 2.6 times (*SD* = 1.3; range: 1–5).

**Table 2.** Subject (*n* = 18) rank, age, number of occurrences as the dominants in dyads (*N* = 47), focal observation time, and rate aggression per hour in focal observation.

| Subjects<br>($n = 18$) | rank | age | number of<br>occurrences as the<br>dominants in dyads | focal observation<br>time (hour) | rate aggression per hour<br>in focal observation |
| --- | --- | --- | --- | --- | --- |
| S127 | 2 | 16 | 2 | 9.66 | 3.83 |
| S112 | 30 | 10 | 5 | 10.00 | 4.60 |
| S86 | 6 | 22 | 2 | 9.98 | 7.72 |
| S85 | 13 | 27 | 4 | 8.66 | 8.89 |
| S67 | 39 | 11 | 3 | 9.99 | 1.70 |
| S36 | 43 | 7 | 3 | 8.32 | 3.00 |
| S25 | 9 | 28 | 1 | 9.94 | 6.14 |
| S20 | 65 | 5 | 1 | 8.32 | 5.17 |
| S2 | 28 | 13 | 2 | 8.99 | 3.78 |
| S53 | 25 | 14 | 1 | 9.97 | 3.21 |
| S136 | 1 | 31 | 3 | 10.00 | 5.40 |
| S138 | 4 | 29 | 4 | 9.99 | 3.30 |
| S108 | 41 | 30 | 1 | 10.00 | 2.30 |
| S80 | 38 | 28 | 1 | 10.00 | 3.20 |
| S82 | 21 | 28 | 4 | 9.33 | 2.79 |
| S7 | 68 | 5 | 4 | 9.89 | 3.94 |
| S54 | 46 | 9 | 2 | 9.99 | 2.30 |
| S47 | 20 | 30 | 4 | 10.00 | 1.20 |
| total |  |  | 47 | 173.03 |  |
| mean | 27.72 | 19.06 | 2.61 | 9.61 | 4.03 |
| SD | 19.84 | 9.58 | 1.30 | 0.59 | 1.96 |

### Rank differences and dominant’s aggressiveness of FID dyads

For the 47 dyads, the mean rank difference was 24.21 (*SD* = 15.77; range: 1–60), indicating that sampling was not strongly biased towards specific rank differences. However, 83% (39 of 47) of these dyads involved individuals with a rank difference of 40 ranks or less, while the maximum possible rank difference was 71. The mean rate of the dominant’s aggression per hour during focal observations was 4.03 (*SD* = 1.96). These rates of aggression were calculated from focal observations and were independently derived from the interactions observed between the 47 dyads. In 83% of subjects (15 of 18 subjects), their aggression rate during focal observations was ≤ 6.00 while maximum aggression rate observed during focal observations was 8.89 times per hour (Figure 1). The repeatability of aggression was *R* = 0.43 (*SE* = 0.19, 95% CI [0, 0.72], *p* = .007). These findings suggest that individual differences in aggression frequency existed and remained relatively consistent throughout the 18-month observation period. However, after controlling for rank, *R* decreased to 0.38 (*SE* = 0.20, 95% CI [0, 0.73], *p* = .097). The correlation coefficient between rank and rate aggression in subjects (*n* = 18) was not significant *r* = −0.34 (*p* = .143).

**Fig 1.**
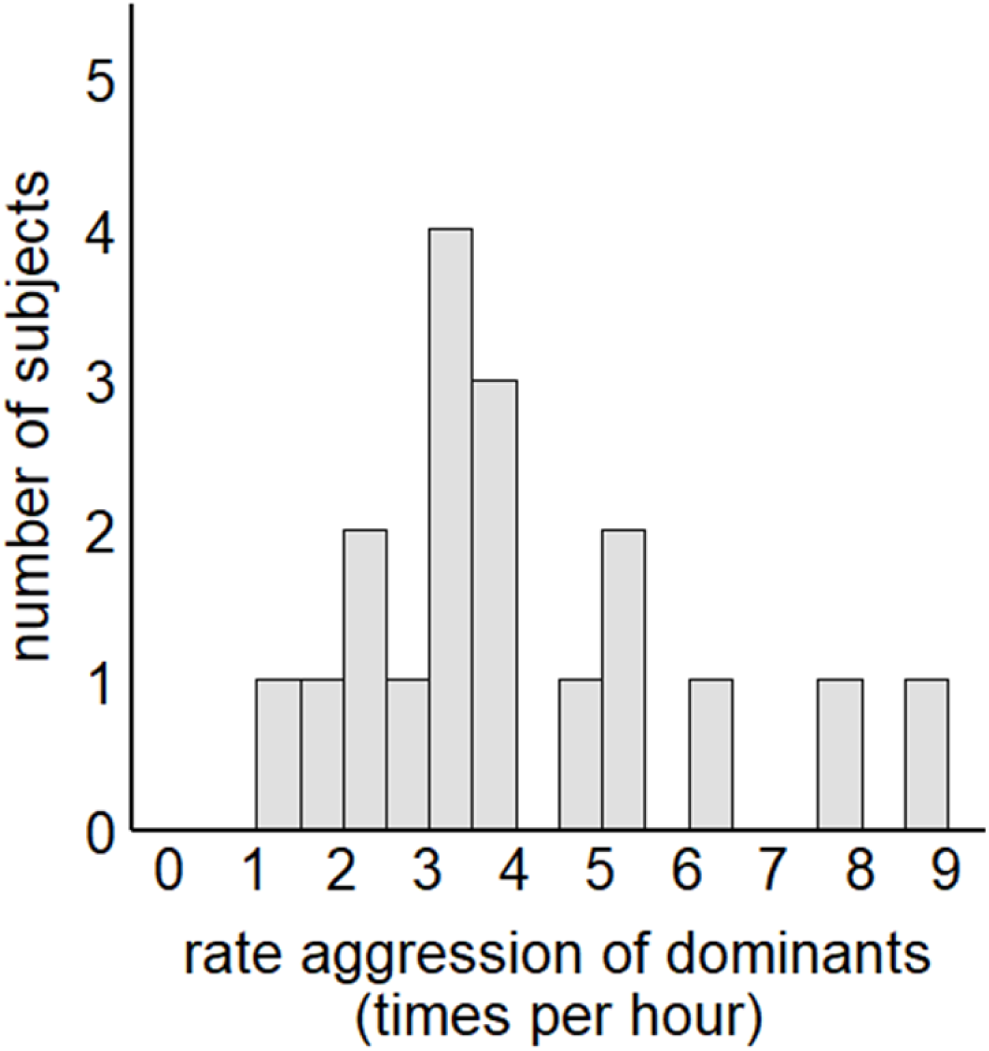
Histograms of the 18 subjects. The y-axis indicates the number of subjects. The x-axis shows rate aggression during focal observations by dominants in FID dyads in increments of 1.0. The focal individuals showed variation in the frequency of aggressive behavior.

### FIDs in relation to rank differences and dominant’s aggressiveness

Of the 47 dyads analyzed, 30 were categorized as ‘long FID’, seven were categorized as ‘short FID,’ and ten were categorized ‘no response.’ Therefore, we combined seven of ‘short FID’ and ten of ‘no response’ to create a total of 17 dyads categorized as “short FID/no response,” which were used in the subsequent analyses. The mean rank difference for the 30 dyads with ‘long FID’ was 25.70 (*SD* = 14.94), while the mean rank difference for the 17 dyads with ‘short FID/no response’ was 21.59 (*SD* = 16.82). The mean rate of the dominant’s aggressive behavior during focal observations in the 30 ‘long FID’ dyads was 4.59 times per hour (*SD* = 2.20). For the 17 ‘short FID/no response’ dyads, the mean rate of aggressive behavior by the dominants was 3.02 times per hour (*SD* = 1.38). The median age of the dominant subjects in the 30 ‘long FID’ dyads was 18.40 years (*SD* = 9.34). For the 17 ‘short FID/no response’ dyads, the mean age of the dominants was 20.35 years (*SD* = 10.47).

**Fig 2.**
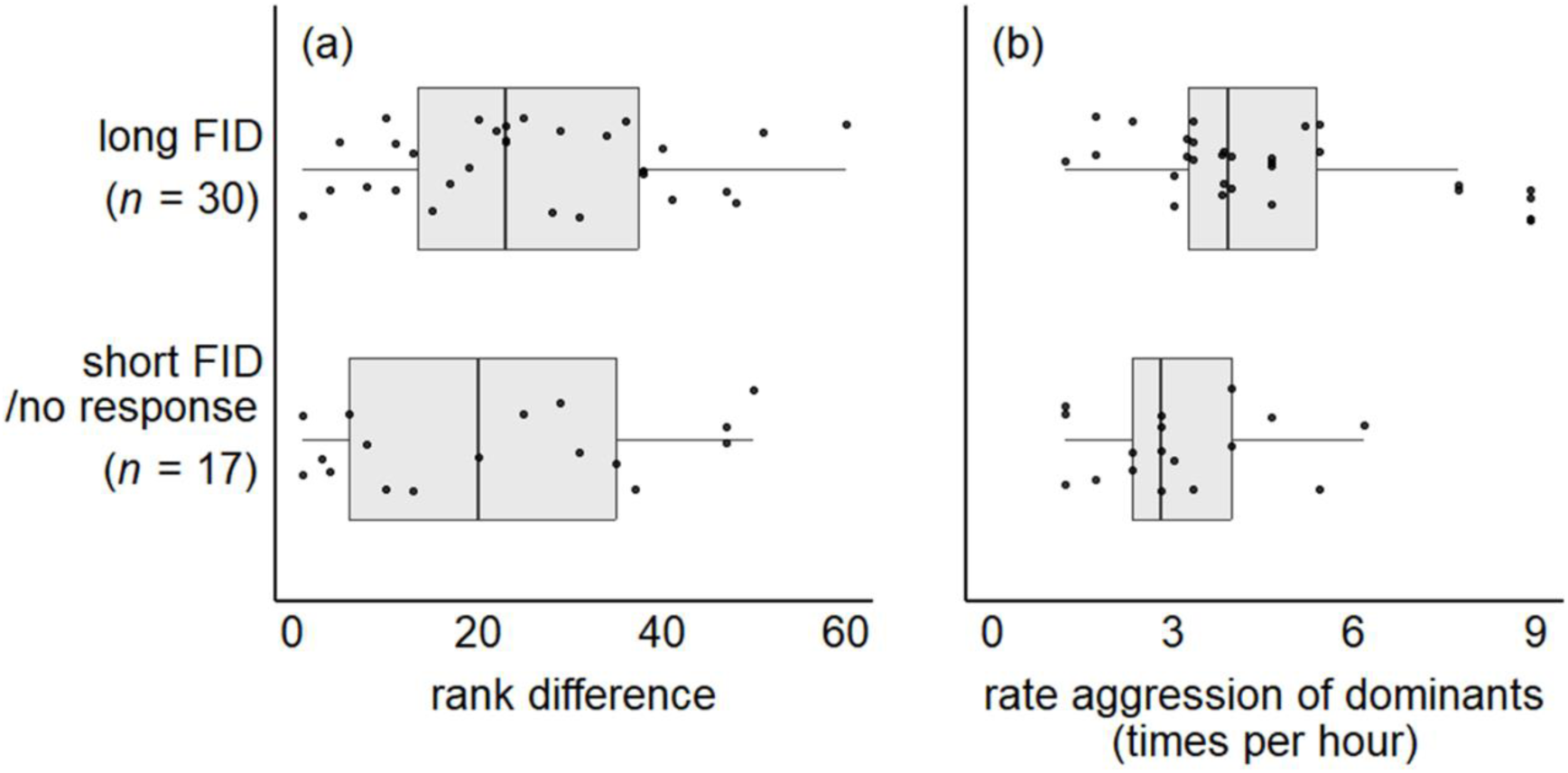
Boxplots of 47 analyzed dyads, comparing those with ‘long FID’ (*n* = 30) and ‘short FID/no response’ (*n* = 17). The x-axes represent (a) rank difference and (b) rate aggression (times per hour) during focal observations by dominants in dyads.

### FIDs in relation to grooming interactions

Among the 47 dyads analyzed, grooming interactions were observed during focal observations in 21% (10 of 47). The proportion of dyads showing a ‘long FID’ was 68% (25 of 37) when no grooming interaction occurred during the focal observation, compared to 50% (5 of 10) when grooming interaction occurred (Fig. 3).

**Fig. 3.**
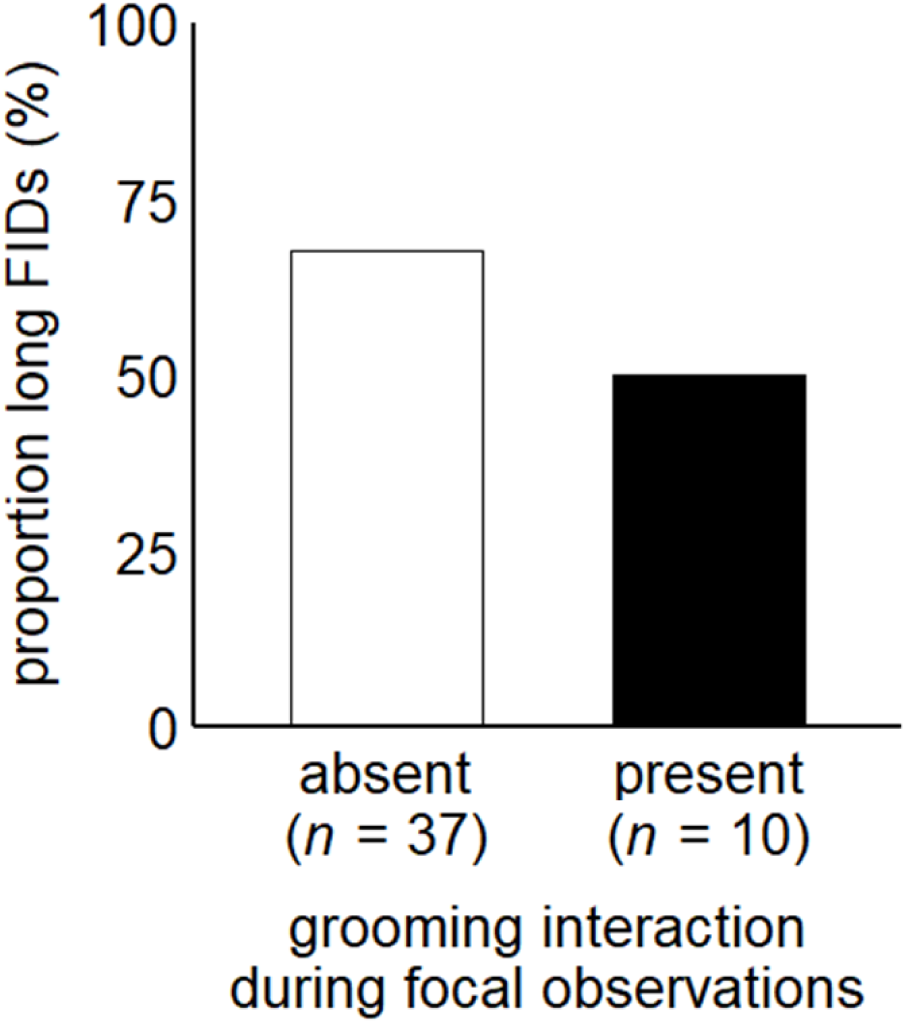
Proportion of the 47 analyzed dyads that exhibited ‘long FID.’ The x-axis represents dyads in which grooming was not observed during focal observations (absent: *n* = 37) and dyads in which grooming was observed (present: *n* = 10).

### FIDs in relation to rank differences and dominant’s aggressiveness

We used a binomial GLM to examine the effects on the likelihood of a ‘long FID’ or ‘short FID/no response’ (Table 3). The analysis revealed that a higher rate of aggression by dominants in FID dyads significantly and positively predicted the likelihood of a long FID (*N* = 47, *Estimate* = 0.47, *SE* = 0.22, *z* = 2.18, *p* = .029). Conversely, rank differences, dominant’s age in FID dyads, and grooming interactions were not significant predictors of FIDs.

**Table 3.** GLM with a binomial error structure for the factors affecting the FIDs (*N* = 47) of subordinates.

| Explanatory variables | Estimate | ± SE | $z$ | $p$ |
| --- | --- | --- | --- | --- |
| (Intercept) | -0.64 | 1.05 | -0.61 | 0.545 |
| rank differences between dyads | 0.02 | 0.03 | 0.76 | 0.445 |
| rate aggression of dominants during focal observation | 0.47 | 0.22 | 2.18 | 0.029 |
| dominant's age in FID dyads | -0.05 | 0.04 | -1.14 | 0.253 |
| grooming interaction (present or absent) | -0.72 | 0.83 | -0.87 | 0.387 |

## Discussion

We revealed that a higher frequency of daily aggressive behaviour by dominant individuals tends to be associated with a longer FID in dyadic supplanting. Neither the rank difference in dyads nor the age of the dominants influenced FID. Furthermore, the occurrence of grooming during focal observations was not related to FID.

Hypothesis 1 was not supported; rank difference did not affect FID. One possible reason for this is that the present study utilized ordinal rank to represent rank difference. Schino & Lasio (2019) reported that even when controlling for ordinal rank, cardinal rank predicted grooming and avoidance relationships. In other words, the frequency of performing dominant behaviours such as supplantings or aggressions, rather than merely a high position in the hierarchy, might better explain interindividual relationships. Had this study been able to assess the difference in the “rates of supplantings” between dominant and subordinate individuals, an effect of rank difference might have been identified.

The second hypothesis was supported: the higher the frequency of aggression by the dominants in daily situations, the longer the FID of the supplanting dyad. This suggests that Japanese macaques may assess how frequently an individual typically exhibits aggressive behaviour, rather than focusing on the rank difference between themselves and a conspecific and subsequently adjust their interindividual distance based on this assessment. Furthermore, this suggests the possibility that monkeys not only identify individuals within their group but also assess their personalities. Additionally, the timid personality of subordinates may have contributed to these results. Regardless of rank difference, timid individuals might have avoided aggressive dominants, potentially obscuring an effect of rank difference. On the other hand, timidity often manifests as anti-predator behaviour, which is less readily observed in provisioned groups. Future research should investigate this aspect by examining measures of an individual’s propensity for anxiety, such as the frequency of scratching.

Contrary to expectations, there was no difference in FID based on whether grooming interactions occurred in focal observations. The small sample size may be a contributing factor to this result. Out of 47 dyads, only 10 engaged in grooming interactions during focal observations. Furthermore, many of the dyadic supplantings were coded as reactions to situations where the dominants “approached” the subordinates. The act of a dominants “approaching” might have readily elicited a flight response from the subordinates, potentially leading to avoidance behaviour even in pairs that are usually affiliative. On the other hand, some studies suggest that prosocial interactions mediated by grooming are based on long-term interactions (Schino & Aureli, 2009; Schino et al., 2007). In the present study, the supplanting events were recorded over a period of three months. If supplantings had been collected over a longer period, the effects of grooming would have become more apparent.

Regarding future directions, most of the dyadic interactions observed in this study occurred around feeding sites. For monkeys, situations involving high-density aggregations with other individuals may increase the necessity to escape from aggressive conspecifics. By examining whether the aggressiveness of dominant individuals lengthens FID in contexts other than feeding, it may be possible to determine if monkeys flexibly use information about conspecific aggressiveness according to the location to regulate their social relationships. Secondly, future studies should record the FID of the same dyad multiple times to verify its consistency. If FIDs are found to be consistent, it would more robustly support the contention that monkeys assess each other’s typical aggressiveness to adjust interindividual distances.

The dyadic data for this study were collected between May and August 2024, whereas the rank difference data were obtained from August 2023 to March 2025. Although dominance rank in Japanese macaques tends to remain relatively stable, some fluctuations may have occurred during this period, potentially influencing the results. Future studies can address this issue by synchronizing the data collection periods for dyadic interactions and rank assessments.

## Acknowledgments

We are grateful for support provided by Mr. S. Asaba, all the staff at Arashiyama Monkey Park. We would like to thank the anonymous referees for their constructive criticism of this manuscript. We thank Y. Nejishima and I. Nakaoka for their insightful comments and supports on the study. This work was supported by JST SPRING, Grant Number JPMJSP2138.

